# Claudin-4 polymerizes after the incorporation of just two extracellular claudin-3 residues

**DOI:** 10.1101/2023.12.15.571691

**Authors:** Rozemarijn E. van der Veen, Jörg Piontek, Marie Bieck, Arbesa Saiti, Hannes Gonschior, Martin Lehmann

## Abstract

Tight junctions play a pivotal role in the functional integrity of the human body by forming barriers crucial for tissue compartmentalization and protecting the body from external threats. Essential components of tight junctions are the transmembrane claudin proteins, which can polymerize into tight junction strands and meshworks. This study delves into the structural determinants of claudin polymerization, utilizing the close homology yet strong difference in polymerization capacity between claudin-3 and claudin-4. Through a combination of sequence alignment and structural modeling, critical residues in the second extracellular segment are pinpointed. Molecular dynamics simulations provide insights into the interactions of and the conformational changes induced by the identified extracellular segment 2 residues, shedding light on the intricacies of claudin polymerization. Live-STED imaging demonstrates that introduction of these residues from claudin-3 into claudin-4 significantly enhances polymerization in non-epithelial cells. In tight junction-deficient epithelial cells, mutated claudin-4 not only influences tight junction morphology but also partially restores barrier function. Understanding the structural basis of claudin polymerization is of paramount importance, as it offers insights into the dynamic nature of tight junctions. This knowledge could be applied to targeted therapeutic interventions, offering insight to repair or prevent barrier defects associated with pathological conditions, or introduce temporary barrier openings during drug delivery.

## Introduction

Barrier formation with the outside world, as well as compartmentalization of tissues within the body are essential to keep the human body functional and protected (1). The necessary barriers are formed by epithelial and endothelial cells, and tight junctions (TJs) seal the paracellular space between them (2). While TJs have several cytosolic and transmembrane components, their backbone is formed by claudins (Cldns) (3). The Cldns contain four transmembrane helices, an intracellular N- and C-terminus and two extracellular segments (ECS1 and ECS2) (4). Through a combination of *cis*- and *trans*-interactions they can polymerize into long TJ strands and even more complex meshworks (5, 6). 25 different Cldns have been described in humans (7, 8). Their expression is tissue-specific, where they provide distinct barrier properties (9). Cldn1, for instance, forms a barrier against water and small molecules in the skin. Its loss in Cldn1-deficient mice leads to postnatal death due to dehydration (10). Alternatively, Cldn16 and Cldn19 form a cation-selective pore in the thick ascending limb of the kidney and their loss causes familial hypomagnesemia with hypercalciuria and nephrocalcinosis (11–13).

While many years of research have advanced our understanding of the structure and function of TJs, they have often been studied as static structures. Nevertheless, it is crucial to understand their assembly and disassembly as well. Not only is proper TJ assembly important for embryonic pattern formation (14), repair of TJ breaks at cell extrusion sites in the mouse intestines also relies on Cldn polymerization (15). Next to TJ (dis)assembly happening naturally, it can also be exploited by pathogens. The C-terminal domain of *Clostridium perfringens* enterotoxin (cCPE) has for instance been shown to disrupt TJ strands through direct interaction with the ECS2 of (mainly) non-junctional Cldns, blocking their assembly (16–19). Moreover, *Entamoeba histolytica* cysteine protease (rEhCP112) has been shown to degrade Cldn1 and Cldn2, thereby increasing TJ permeability (20). A better understanding of TJ (dis)assembly could help us prevent or compensate for barrier defects due to these pathogens. Conversely, this knowledge could also be used in drug delivery, where temporary TJ opening could improve delivery efficiency to certain tissues, especially across the very tight blood-brain barrier (21).

Many studies have concentrated on signaling pathways and scaffolding proteins affecting tight junction formation (22–24). However, a more direct approach to modifying tight junction (dis)assembly would be to influence Cldn polymerization. Based on the Cldn15 crystal structure (4) models of Cldn15 polymer-based TJ strands have been suggested (6, 25–27). More recently, polymeric Cldn10b models have been added to this (28, 29), but models for other Cldns or combinations of Cldns are still missing. From these models, combined with experimental data, it has been proposed that Cldns assemble into *cis*-oligomers before *trans*-interactions at cell-cell contacts trigger polymerization (28, 30, 31). Furthermore, the Cldn ECS2 was found to mediate the *trans*-interaction and thereby drive polymer assembly (28, 32).

Further understanding of Cldn polymerization has been gained from recent imaging advances. Through pulse-chase labeling in kidney epithelial MDCKII cells, it was shown that newly synthesized Cldns are incorporated into the TJ from the basal side (33). Moreover, structured illumination microscopy in living Cldn2-expressing Rat-1 fibroblasts demonstrated additional incorporation at strand break sites (34). Non-epithelial cells like fibroblasts have been used since a long time to understand Cldn polymerization, since expression of Cldns in these cells can reconstitute TJ strands (35). In a recent study in COS-7 cells, stimulated emission depletion (STED) microscopy revealed that only half of the Cldn family can form polymers on their own (36). Interestingly, in this study Cldn3 polymerized, but its close homolog Cldn4 only gave diffuse membrane staining. Another study in U2OS cells also demonstrated the inability of Cldn4 to form a meshwork (37). Finally, in L-fibroblasts, Cldn3 was found to enrich at the cell-cell contact and form a TJ-like meshwork (5, 16). Cldn4, however did not show a similar enrichment (16).

In this study we therefore decided to compare Cldn3 and Cldn4 and explore the structural determinants of TJ polymer formation. Given the close homology but strong difference in polymerization capacity of these two proteins, we saw a unique opportunity to understand TJ assembly in more detail. Utilizing sequence alignment and structural modeling, we pinpoint two ECS2 residues that could account for the disparity in polymerization of Cldn3 and Cldn4. Introduction of the corresponding Cldn3 residues in Cldn4 induces meshwork formation, as evidenced by live-STED imaging in COS-7 cells. Moreover, it can partially restore the barrier function and morphology of TJs in TJ-deficient MDCKII cells. We thus discovered an intramolecular motif that shapes the polymerization properties of Cldns.

## Results

### Structural comparison of Cldn3 and Cldn4 indicates residues in ECS2 as critical determinants for polymerization

In line with a previous study using YFP-tagged proteins in COS-7 cells (36), we found that human SNAP-tagged Cldn3 and Cldn4 are, respectively, able and unable to polymerize in this model system [Fig. 1*A*]. To determine which residues could underlie this difference, we therefore compared their sequences in a multiple sequence alignment. Cldn6, Cldn8, Cldn9 and Cldn17 were included in the alignment, as they are evolutionary closely related to Cldn3 and Cldn4 [Fig. 1*B*]. Cldn6 and Cldn9 can form homopolymers in COS-7 cells, whereas Cldn8 and Cldn17 cannot (36). This allowed us to look for amino acid differences that were conserved among meshwork versus non-meshwork formers. In this comparison [Fig. S1], we identified a key difference in ECS2: E152 in Cldn3 versus S153 in Cldn4. The larger, negatively charged glutamic acid in Cldn3 is conserved among Cldn6 (E153) and Cldn9 (E153), whereas Cldn4, Cldn8 (V154) and Cldn17 (I154) all have smaller, polar/hydrophobic residues at this site.

**Figure 1.**
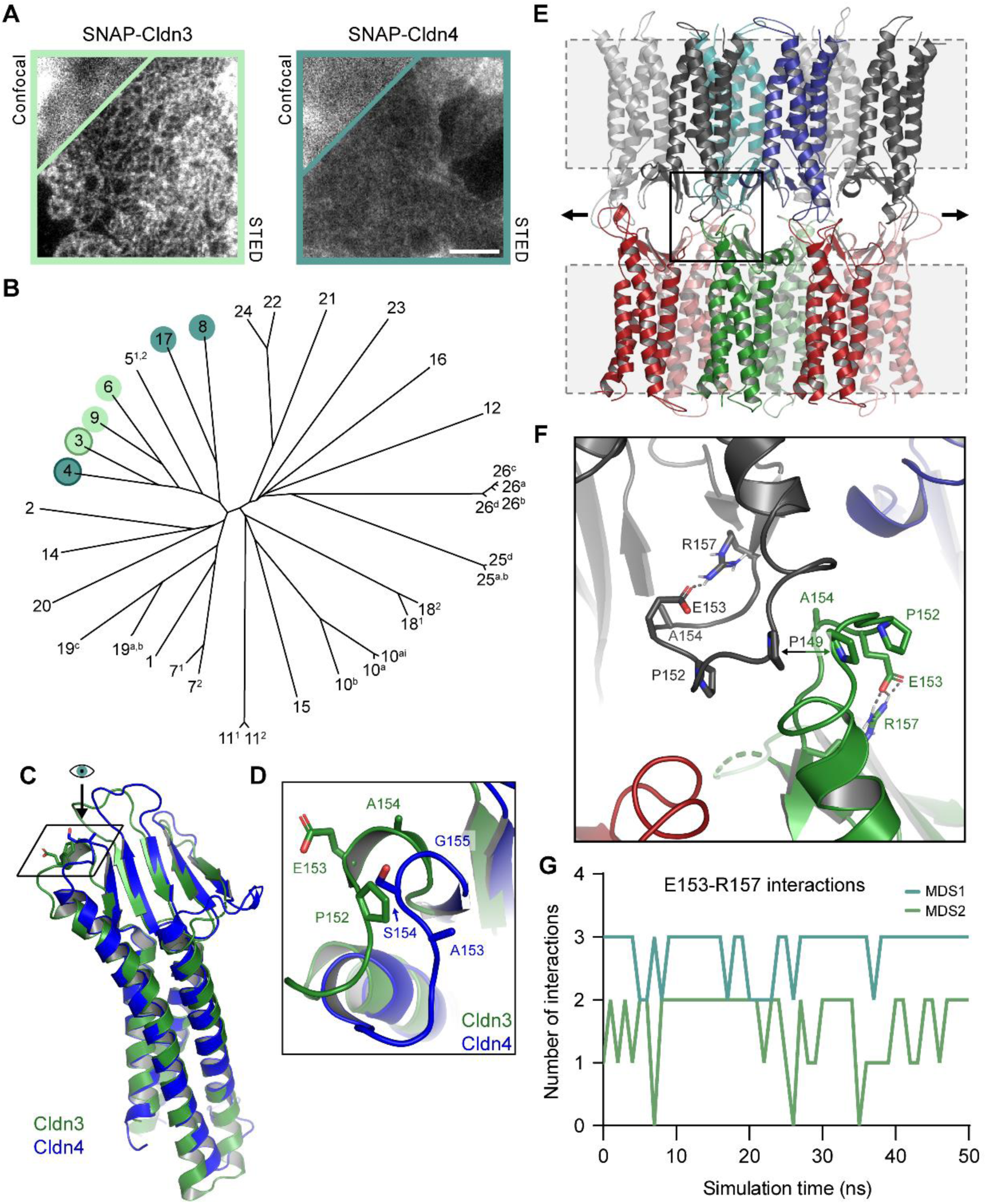
Sequence alignment and structural investigation of Cldn3 and Cldn4 points towards two residues in ECS2 that may be critical for polymerization. *A*, Representative STED and confocal (insert) images of SNAP-Cldn3 and SNAP-Cldn4 in COS-7 cell-cell overlaps. Scale bar: 1 µm. *B*, Phylogenetic tree of the Cldn family, following the nomenclature suggested by Mineta et al. (7), demonstrating that meshwork-forming Cldn3, Cldn6 and Cldn9 (in green), and non-meshwork-forming Cldn4, Cldn8 and Cldn17 (in blue) fall into the same cluster. *C*, Aligned crystal structures of mouse Cldn3 (green) and human Cldn4 (blue) monomers, derived from cCPE-claudin complexes (PDB 6AKE (42), 7KP4 (43)). *D*, Top view of ECS2 region indicated in (*C*), demonstrating key residue differences (O atoms in red) between Cldn3 (green: P152, E153, A154) and Cldn4 (blue: A153, S154, G155) that result in distinct backbone conformations. *E*, Cldn3 dodecamer model based on previously generated Cldn10b model (29). Snapshot of the MD simulation, showing the overall conformation of all twelve Cldn3 subunits. Cell membranes (not included in simulation) are depicted as gray dashed boxes. Arrows indicate the direction of strand elongation. *F*, Close-up of region outlined in (*E*) showing trans-interacting ECS2s of two Cldn3 subunits (black and green). Intra-molecular electrostatic interactions between E153 and R157 (dashed lines; O, red; N, blue) shape the ECS2 turn together with the rigid P152, allowing the P149 residues of the two subunits to get in close trans-proximity. *G*, Number of E153-R157 interactions in four central subunits of the Cldn3 dodecamer during two 50 ns MD simulations.

Crystal structures of mouse Cldn3 and Cldn4 (complexed with the C-terminal domain of *Clostridium perfringens* enterotoxin; cCPE) allowed us to examine their ECS2s in more detail [Fig. 1*C* and *D*]. A comparison showed us that Cldn3 and Cldn4 overall have a very similar structure. Interestingly, the residue identified in our alignment is however found in a region with conformational differences, not only caused by this residue (E153/S154; E152/S153 in human), but also the neighboring residues (P152/A153 and A154/G155). However, in both Cldns, the residues were pointing outwards from the ECS, in the direction that a *trans*-interacting subunit would be positioned (6, 28) [Fig. 1*D*].

To understand the importance of this region in polymer formation, we created an oligomeric Cldn3 homology model, based on experimentally derived constraints (28) and a Cldn10b dodecamer template (29). The model was minimized and equilibrated using molecular dynamics (MD) simulations with water as solvent and the transmembrane helices fixed by constraints [Fig. 1*E*]. As opposed to it facing outwards, as it is in the Cldn3 crystal structure, E153 was found to point inwards in the dodecamer model [Fig. 1*F*]. Here it frequently electrostatically interacts with R157 [Fig. 1*G*], thereby shaping the conformation of the ECS2 turn. This conformation is further stabilized by the rigid, neighboring P152 and favors close proximity of the P149 residues of two trans-facing subunits [Fig. 1*F*]. Based on these simulations it is thus suggested that the combined rigidity and electrostatic interaction in the ECS2 of Cldn3 (both missing in Cldn4) contribute to a stronger trans-interaction and polymer formation.

### Substituting two Cldn4 residues in the ECS2 with their Cldn3 homologs induces meshwork formation in living non-epithelial cells

In order to test whether the combined rigidity and negative charge are sufficient to promote meshwork formation, we introduced these residues in Cldn4 (A153P and S154E, from now on denoted as Cldn4^ECS2-mut^). This mutant was analyzed together with Cldn3 and Cldn4 in living COS-7 cells, in order to rule out any artifacts or loss of meshworks due to fixation. In contrast to our previous report using fixated COS-7 cells (36), Cldn4 was occasionally capable of forming meshworks in these living cells [Fig. 2*A*, Video 1]. We analyzed a large amount of cell-cell overlaps and classified whether a dense meshwork, a thin meshwork, isolated strands or membrane staining was present [Fig. 2*B* and *C*]. In this way we could validate that Cldn3 easily polymerizes, forming meshworks 74% of the time, whereas Cldn4 predominantly demonstrates diffuse membrane staining (78%). Interestingly, the two point mutations in the ECS2 of Cldn4 strongly induced polymerization; a meshwork was formed in 57.5% of the cases.

**Figure 2.**
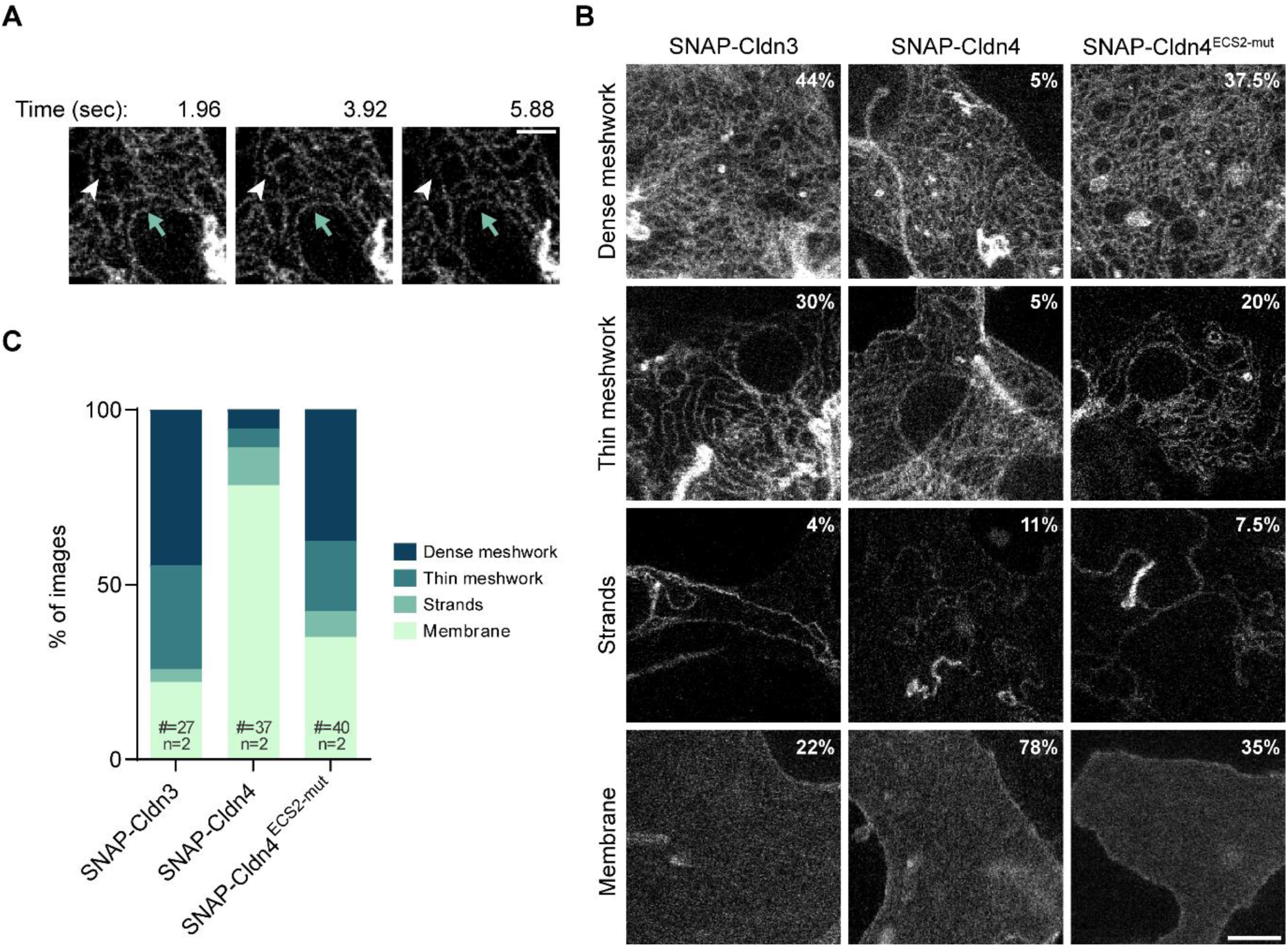
Systematic STED imaging and classification in living COS-7 cells demonstrates that altering two ECS2 residues of Cldn4 to match those found in Cldn3 can effectively rescue Cldn4’s ability to polymerize. *A*, Snapshots of a SNAP-Cldn4 meshwork in a COS-7 cell-cell overlap, showing upwards meshwork movement (blue arrow) and strand remodeling (white arrowhead). A Guassian blur of 10 nm was applied to improve image clarity. Scale bar: 500 nm. *B*, Representative images of different structures formed by SNAP-Cldns in COS-7 cell-cell overlaps, along with the corresponding prevalence percentages, pooled from two independent experiments. Scale bar: 1 µm. *C*, The occurrence of different (non)-polymerized structures in overlaps between COS-7 cells upon SNAP-Cldn expression, pooled from two independent experiments.

### Introduction of ECS2-mutated Cldn4 partially rescues TJ morphology and barrier function in epithelial cells

Given the compelling polymerization rescue exhibited in non-epithelial cells, we proceeded to investigate how Cldn4^ECS2-mut^ affects the TJ properties in epithelial cells. For this, we made use of a TJ-deficient MDCK cell line, derived by knockout of Cldn1, Cldn2, Cldn3, Cldn4 and Cldn7, known as Claudin quintuple knockout (qKO) cells (38). Using a retroviral expression system, FLAG-tagged Cldn3, Cldn4 and Cldn4^ECS2-mut^ were introduced in qKO cells at similar expression levels [Fig. 3*A*]. All proteins were expressed above their endogenous levels in MDCKII cells [Fig. 3*B* and *C*] and localized to the plasma membrane [Fig. 3*D* and *E*]. Except for slight cell to cell variability, protein expression was homogeneous and found in all cells [Fig. 3*D*]. Interestingly, all proteins were found both in the TJ and in the lateral cell membrane [Fig. 3*E*]. When comparing the overlap with the TJ marker ZO1, Cldn3 exhibited the highest, Cldn4 the lowest, and Cldn4^ECS2-mut^ an intermediate degree of overlap.

**Figure 3.**
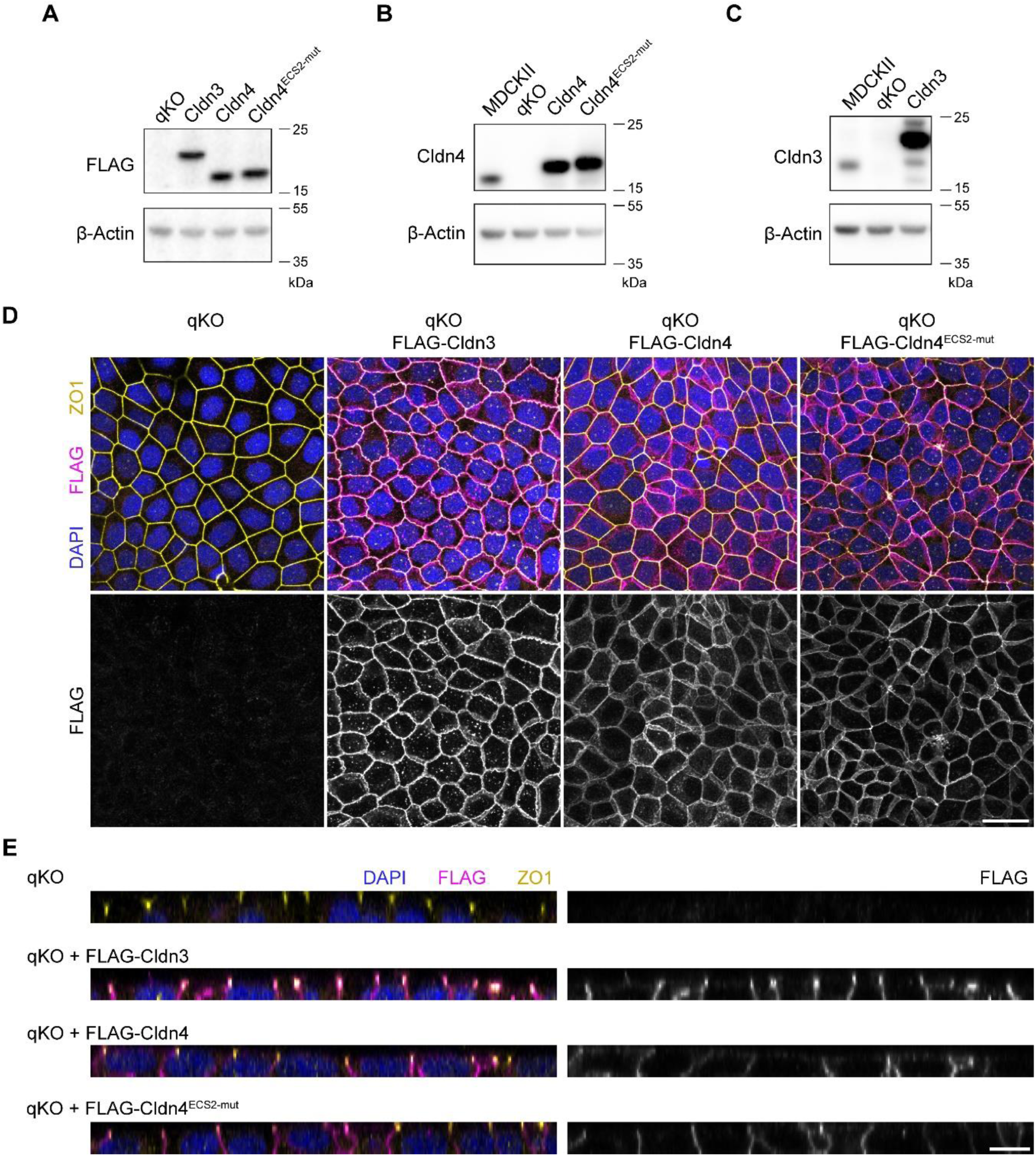
Establishment of stable Cldn3, Cldn4 and Cldn4^ECS2-mut^ expression in TJ-deficient epithelial cells. *A-C*, Stable expression of FLAG-Cldn3, -Cldn4 and -Cldn4^ECS2-mut^ in qKO cells was validated using immunoblotting against the FLAG-tag (*A*), Cldn4 (*B*), Cldn3 (*C*) and β-actin (loading control). *D-E*, Membrane and TJ localization of FLAG-Cldn3, -Cldn4 and -Cldn4^ECS2-mut^ (magenta/grey) in qKO cells, co-stained with DAPI (blue) and ZO1 (yellow) as a TJ marker. Representative maximum intensity projections (*D*) and orthogonal views (*E*) of an 8 µm z-stack are shown. Scale bars: 25 µm (*D*) and 10 µm (*E*).

When assessing the ZO1 staining in more detail, clear differences in junctional morphology were observed depending on the expressed Cldn [Fig. 4*A*]. qKO cells display a bright, linear ZO1 signal, as reported before (38, 39), which is unaltered by Cldn4 expression in these cells. Contrastingly, Cldn3 expression led to a more ruffled and dimmer ZO1 signal, similar as observed in MDCKII cells expressing all endogenous Cldns (40). Cldn4^ECS2-mut^ expression also reduced ZO1 signal and introduced ruffles, though not as strongly as Cldn3. Automatic quantification of the zigzag indexes in the different cell types [Fig. 4*B* and *C*] produced similar values for qKO cells and cells expressing Cldn4 [Fig. 4*D*]. Cldn3-expressing cells had the highest index and the indexes of Cldn4^ECS2-mut^-expressing cells fell in between, but were significantly different from those found for Cldn4. These values correlate well with what was found upon ZO1 knockout and reconstitution in MDCKII cells (40): qKO and Cldn4 cells have a similar lack of ruffles as ZO1 KO cells and Cldn3 introduces ruffles comparable to low-level ZO1 reconstitution.

**Figure 4.**
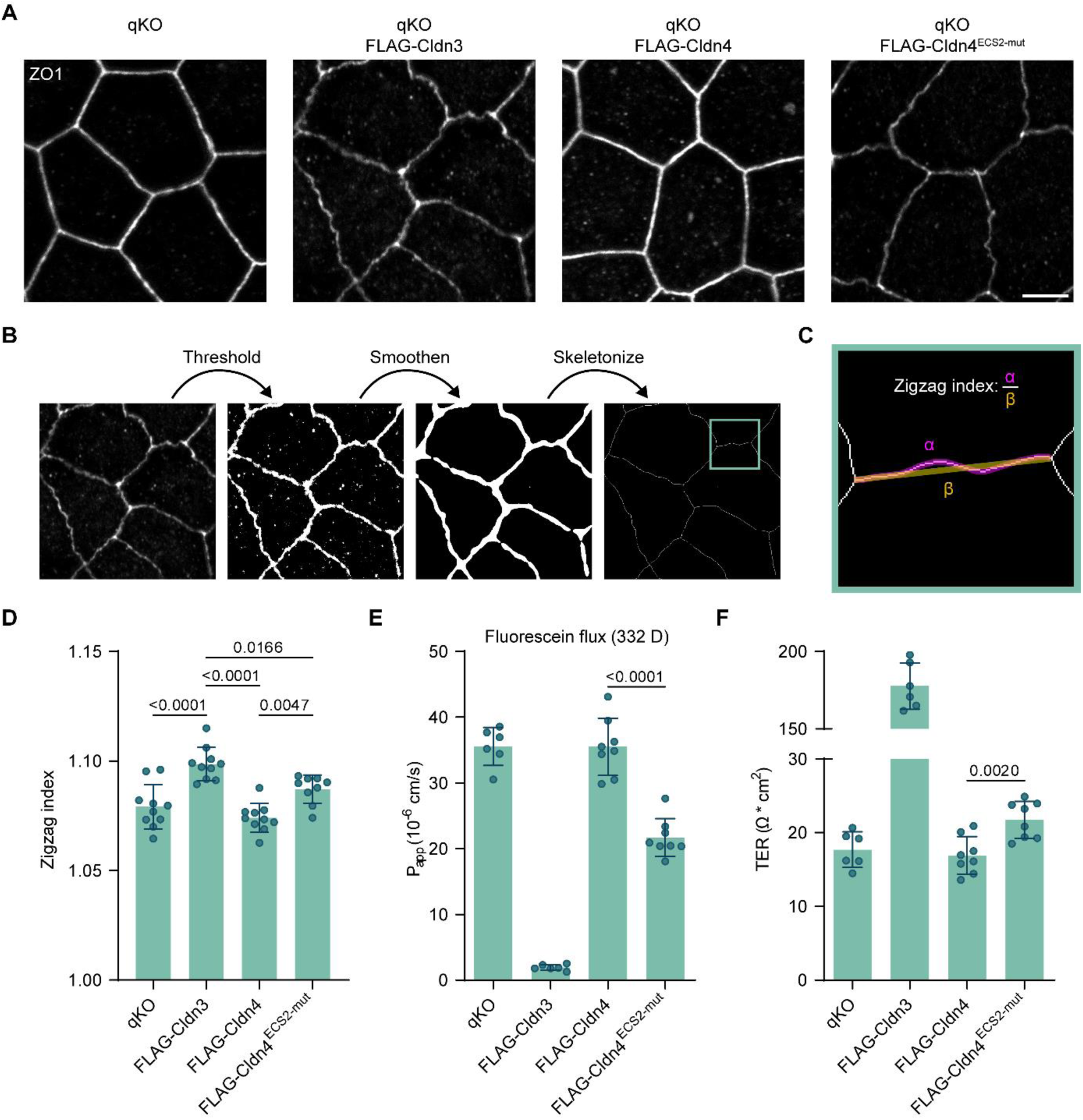
Cldn4^ECS2-mut^ expression can partially rescue the formation of a ruffled TJ and a barrier against small molecules and electrolytes in qKO cells. *A*, ZO1 staining in qKO cells (expressing FLAG-Cldn3, -Cldn4 or Cldn4^ECS2-mut^), demonstrating distinct TJ morphologies. Scale bar: 5 µm. *B-C*, Description of the automated pipeline used for quantification of the zigzag index of the TJs. On each image the same threshold was applied. Afterwards the images were smoothened and skeletonized (*B*). For each cell-cell contact spanning from one tricellular junction to another, the distance covered by the ZO1 signal (α) as well as the shortest distance between the two tricellular junctions (β) was determined. Division of these two resulted in the zigzag index, which was determined as an average per image (*C*). *D*, ZO1 zigzag indexes quantified as described in (*B-C*) for qKO cells (expressing FLAG-Cldn3, -Cldn4 or Cldn4^ECS2-mut^). Each dot represents the average zigzag index from one image (24-71 cell-cell junctions were analyzed per image, n=9-10). Mean ± SEM. One-way ANOVA with Tukey’s multiple comparison test. Adjusted p-values ≤ 0.05 are shown. *E*, Paracellular permeability for fluorescein (332 D) across qKO cell monolayers (expressing FLAG-Cldn3, -Cldn4 or Cldn4^ECS2-mut^). Every dot represents one transwell filter (n=6-8). Mean ± SEM. Two-tailed Student’s *t* test. *F*, Evaluation of transepithelial electrical resistance (TER) of qKO cell monolayers (that expressed FLAG-Cldn3, -Cldn4, or -Cldn4^ECS2-mut^). Each dot describes one transwell filter (n=6-8). Mean ± SEM. Two-tailed Student’s *t* test.

Finally, we evaluated whether reconstitution of the different Cldns in qKO cells could improve TJ barrier function. Due to the absence of TJ strands, small molecules and electrolytes can easily permeate between qKO cells (38). However, it has been shown that reconstitution of barrier Cldns like Cldn3 can rescue these barrier deficiencies (36, 41). As done in these past studies, we measured the permeability of a small fluorescent tracer molecule (fluorescein, 332 D) [Fig. 4*E*], as well as the transepithelial electrical resistance (TER) [Fig. 4*F*]. We were able to reproduce the barrier deficiency in qKO cells and barrier rescue in Cldn3-expressing cells. Surprisingly, the reintroduction of Cldn4 had no positive impact on barrier function, with no observable changes in fluorescein flux or TER. In contrast, Cldn4^ECS2-mut^ did show the ability to restore barrier function, evident by a marked reduction in fluorescein flux and a slight increase in TER. Overall, we show that Cldn4^ECS2-mut^ can not only rescue polymerization in non-epithelial cells, but also TJ properties in epithelial cells.

## Discussion

In this study we identified a small structural motif with a significant effect on Cldn polymerization capacity. Introduction of only two Cldn3 amino acid residues into the ECS2 of Cldn4 (A153P, S154E) still leaves 19% difference in their protein sequence, yet largely rescues meshwork formation in COS-7 cells. We postulate, based on homology models and MD simulations, that these residues induce an ECS2 conformation favoring *trans*-interaction through rigidity (of the proline) and an intramolecular salt bridge (of the glutamic acid with R157/158 in Cldn3/4). Given the importance of *trans*-interactions for polymerization (28–30), this conformational difference can heavily impact polymer formation. When retrovirally introduced into TJ-deficient MDCKII (qKO) cells, it is evident that FLAG-Cldn4 fails to form a barrier for electrolytes and small molecules. Contrary, its mutated counterpart successfully establishes a (partial) barrier and alters junctional morphology from a straight to a ruffled pattern. Additionally, we illustrate that live-STED imaging is an effective tool for identifying morphological distinctions among meshworks formed by various Cldns, thus eliminating potential impacts of fixative agents. This adds to past studies in which super-resolution microscopy was used, either for comparing Cldn meshwork morphology in fixed samples (36) or for analyzing dynamics of only one Cldn (34).

The emphasis of our research was on intrinsic structural properties driving Cldn polymerization, but it should be noted that polymerization is also influenced by context. This context can for instance be the level of expression. In MDCKII cells, Cldn4 overexpression increases the TER, decreases the flux of cations and creates a larger TJ meshwork (44, 45). Its KO on the other hand has not apparent effect on the TER or on the ion and small molecule permeability (46). Similar observation were made in opossum kidney cells: overexpression increases the TER, KD does not (47). The presence of other Cldns can also make a difference. While Cldn4 in COS-7 cells does not polymerize on its own, it has been shown to copolymerize with Cldn3 and Cldn8 (36). The latter has been suggested to induce formation of a chloride channel in the collecting duct (48), which would explain why in collecting duct cell lines, Cldn4 KD does not decrease but increases the TER (48). On the other hand, Cldn4 itself seems to disrupt polymerization by cation channel-forming claudins like Cldn2 and Cldn15 and thereby can reduce cation flux (37).

Other factors that could influence Cldn4 polymerization are the different N- or C-terminal tags used in the studies and the species origin of the protein. The importance of all of these factors is especially evident when comparing our data to a recent study by Furuse et al (41). In the same qKO cell line, they find that Cldn4, which did not create a barrier in our experiments, makes TJ strands and introduces a strong barrier against solutes. Despite sharing a similar experimental set-up, the two approaches diverge in various aspects. They employ clones that exhibit a significant overexpression of untagged dog Cldn4, whereas we use a mixed cell population expressing N-terminal FLAG-tagged human Cldn4 at a comparatively lower expression level.

While mutating Cldn4 strongly rescued polymerization in COS-7 cells, the effect on the TJ barrier in the qKO cells was smaller. This discrepancy may result from the difference in cell type: COS-7 cells lack the polarity of epithelial cells. Therefore, they may depend less on Cldn trafficking, whereas in the MDCKII-derived qKO cells the proteins need to be correctly targeted to the TJ. This could especially be important for Cldn4, which was shown to have high TJ turnover (compared to Cldn2) in MDCKII cells (33). Upon expression in qKO cells, all Cldns in this study did not only show up in the TJ, but also in the lateral membrane. Endogenous Cldn3/4 and Cldn4 with an N-terminal SNAP-tag have also been shown to localize laterally in MDCKII cells (33). We found TJ localization to be strong for Cldn3, weak for Cldn4 and intermediate for Cldn4^ECS2-mut^. TJ targeting most likely not only depends on the ECSs, but also on other Cldn domains, in particular the cytosolic C-terminal region. Cldn4 has for instance been shown to depend on its C-terminal region for membrane trafficking (33). The C-terminus of both Cldns has a PDZ motif for binding to the cytosolic adaptor ZO1 (49, 50). Without ZO1 and ZO2, Cldn3 moves from the TJ to the lateral membrane in EpH4 cells (51). Given that Cldn3 and Cldn4 have a vastly different C-terminal region [Fig. S1] that could result in different regulation of ZO binding, their trafficking in epithelial may be very different. Next to this, trafficking may be affected by the absence of other Cldns. TJ targeting of Cldn3 and Cldn4 was, for instance, shown to depend on Cldn23 in intestinal epithelial cells (52). Similarly, it has been suggested that Cldn8 can improve the Cldn4 recruitment to the TJ (48). On the contrary, Cldn2 seems to inhibit Cldn4 TJ localization, as its KO enhance Cldn4 signal at the TJ (33).

A ruffled junctional morphology as we observed in qKO cells upon expression of only Cldn3 or Cldn4^ECS2-mut^ has been reported in a wide array of papers (39). Usually, ruffle formation is associated with TJ-regulating conditions like hypoxia or anoxia. Here ruffling seems to increase paracellular permeability by increasing TJ circumference (53–55). This is in stark contrast to our findings, since a loss of ruffling in our system correlates with an increase in paracellular permeability. However, it has been shown that ruffling depends on Cldns (38), ZO1 (40), and their link to the actin cytoskeleton (39, 56), as the loss of any of these leads to a straight, very leaky TJ. We therefore postulate that the combination of a very straight and permeable TJ in our system implies the absence of polymerized Cldns, whereas the ruffles imply that polymerization has been rescued.

The Cldn3 polymer model created in this study was based on previously generated Cldn15 and Cldn10b models (6, 29). Recently, another group published computational models for Cldn3 and Cldn4 (52). Nevertheless, their study did not cover several subunit interfaces involved in polymerization, including the ECS2-cotaining interfaces which were the focus of this study. Our modeling data suggest that for Cldn3 a E153-R157 interaction and the presence of the rigid P152, both absent in Cldn4, shape the ECS2 conformation and thereby strengthen ECS2-ECS2 *trans*-interaction. Of note, the ECS2 conformation and E153 orientation in the simulated Cldn3 oligomers differs from the ones observed in Cldn3 and Cldn4 crystal structures. However, in the latter the ECS2 conformation is strongly influenced by binding of cCPE resulting in the crystalized claudin-cCPE complexes (42, 43). It is likely that the Cldn ECS2 can adopt different conformations: one stabilized by cCPE binding, the other by Cldn-Cldn *trans*-interaction. Similarly, conformational differences in the TM3/ECS2 of Cldns were reported to affect Cldn *cis*-interaction (42).

Interestingly, the electrostatically fitting glutamic acid-arginine pair is conserved among barrier-forming Cldn3, Cldn6 and Cldn9. Moreover, other barrier-forming claudins have residues at both positions that fit by hydrophobicity (Cldn1, Cldn5, Cldn7, Cldn14 and Cldn19). Channel forming Cldn2, Cldn10b and Cldn17 are charged only at one of both positions (3) and in Cldn10b channel models, E153 points towards the pore center, where it contributes to cation guidance (29). This covariation suggests that the positions corresponding to E153 and R157 in Cldn3 are in several claudins involved in barrier regulation. At least for R157 in Cldn3 and the corresponding Y158 in Cldn5 involvement in strand formation has been shown previously (32, 57). Other motifs involved in claudin polymerization have been identified previously (3, 28, 29). However, the corresponding sequences are highly conserved, also between Cldn3 and Cldn4. These motifs could therefore not explain the different polymerization capacity of the two Cldns, in contrast to the novel motif identified in this study. Detailed MD simulations of membrane-embedded oligomers of Cldn3, other barrier-forming claudins and Cldn4 are expected to provided further mechanistic information.

Cldn3 and Cldn4 were the first receptors discovered for CPE, the enterotoxin of *Clostridium perfringens* (58). This bacterial toxin causes the symptoms associated with food poisoning in roughly one million United States citizens each year (59, 60). Even though Cldn3 and Cldn4 are both high affinity CPE receptors, Cldn4 binds the toxin with higher affinity (61). In a study investigating the mechanism of cCPE-Cldn interaction, a similar Cldn3-resembling amino acid substitution in Cldn4 ECS2 (Cldn4 A153P, S154E, G155A) reduced cCPE binding by roughly 22% (62). Modeling and experiments have implied that CPE only binds to non-junctional Cldns, as TJ Cldns would be sterically inaccessible (17–19). Thus, we postulate that the identified ECS2 motif not only provides Cldn4 with a direct advantage over Cldn3 in CPE binding, but also an indirect advantage. Due to its inhibitory effect on polymerization, Cldn4 would be a more accessible non-junctional CPE receptor.

In conclusion, we report the identification of a structural determinant of Cldn polymerization and give insight in the (importance of the) ECS2 interface in Cldn polymers. This knowledge can form the basis for further systematic mutational studies and modeling efforts to understand polymer formation across the Cldn family. Ultimately, this could support the design of innovative drugs targeted at precisely modulating the tight junction barrier in different tissues.

## Experimental procedures

### Constructs and molecular cloning

All Cldn sequences used in this study were human. The SNAP-Cldn3 and SNAP-Cldn4 constructs used in this study have been published in a past study (36). The A153P, S154E mutations in ECS2 of SNAP-Cldn4 were introduced using two primers: 5’-AATCCGCTGGTGCCGGAAGGGCAGAAGC-3’ and 5’- GTAGAAGTCTTGGATGATGTTGTGGGCCGTCCAG-3’ (BioTeZ Berlin-Buch GmbH). Primers were 5’- phosphorylated and site-directed mutagenesis through two-step PCR amplification and ligation was executed according to manufacturer’s protocol (Thermo Scientific Phusion Site-Directed Mutagenesis Kit, F541). Ligation products were used to heat shock transform self-made chemically competent TOP10 *E. coli* cells. Selected colonies were cultured up and DNA was isolated from them (Macherey-Nagel, 740420.50). Successful creation of SNAP-Cldn4^ECS2-mut^ was verified with sequencing (LGC Genomics GmbH, Berlin).

pLIB-CMV-FLAG-Cldn4-Puro and pLIB-CMV-FLAG-Cldn4^ECS2-mut^-Puro were created from the previously published pYFP-Cldn4 (36), the above described SNAP-Cldn4^ECS2-mut^ and the published pLIB-CMV-GFP-Puro construct (36). FLAG-tagged Cldn4 was PCR amplified with the following primers: FW 5’- TATAACCGGTATGGATTACAAGGATGACGACGATAAGCTGTACAAAAGCTTGGTACCGAGCTCGGATCCAATGG-3’ and RV 5’-TATAGCGGCCGCGGATTATCTTACACGTAGTTGCTGGC-3’. Amplification of FLAG-tagged Cldn4^ECS2-mut^ was done with 5’- TATAACCGGTATGGATTACAAGGATGACGACGATAAGCTGTACAAAAGCTTGGTACCGAGCTCGGATCCAATGGCCTCCATG-3’ and the same RV primer as Cldn4. Two-step PCR amplification was done according to manufacturer’s protocol (Thermo Fisher Phusion High-Fidelity DNA Polymerase, F530S) and followed with a PCR clean-up (Macherey-Nagel, F40609.250). PCR products and plasmids were restricted with FastDigest BshTI (Thermo Fisher, FD1464) and NotI (Thermo Fisher, FD0593) enzymes. Correct products were isolated with gel electrophoresis and gel extraction (Macherey-Nagel, F40609.250), and were ligated with T4 DNA ligase (Thermo Fisher, EL0016). HB101 cells (Promega, L2011) were heat shock transformed, DNA was isolated (Macherey-Nagel, 740420.50) and the sequences were checked with sequencing (LGC Genomics GmbH, Berlin). pLIB-CMV-FLAG-Cldn3-Puro, as well as the pMD2.G retroviral envelope and pCIG3.NB packaging plasmids have been published before (36).

### Cell culture, transient transfection and SNAP-labeling

In this study COS-7 (ATCC CRL 1651), HEK293T (ATC CRL 11268), MDCKII (ECACC 00062107) and MDCKII claudin quintuple knockout (qKO, courtesy of Prof. Mikio Furuse, NIPS, Okazaki Japan) cells were used. All were cultured at 37 °C and in a 5% CO_2_ atmosphere in high glucose DMEM (Thermo Fisher, 11965084), supplemented with 100 μg/ml penicillin-streptomycin (Gibco, Thermo Fisher, 15140122) and 10% FBS (v/v) (Gibco, Thermo Fisher, 10082147). qKO cells stably expressing FLAG-Cldns were maintained with the addition of 2 µg/mL puromycin (Thermo Fisher, A1113803) in the medium.

For fixed STED imaging, 15.000 cells were seeded per well of a µ-Slide 8-well glass bottom dish (Ibidi, 80827). For live STED 200.000 cells were seeded in a glass-bottom 35 mm µ-dish (Ibidi, 81156). Both dishes were previously coated with 2% Cultrex Reduced Growth Factor Basement Membrane Extract (v/v) (Bio-techne, 353600502) in medium. 24 hours after seeding, cells were transfected using Lipofectamine 2000 Transfection Reagent (Thermo Fisher, 11668019), according to manufacturer’s protocol. After 6 hours the medium containing transfection reagents was replaced by FluoroBrite DMEM medium (Thermo Fisher, A1896701), supplemented with 10% (v/v) FBS (Gibco, Thermo Fisher, 10082147). Another 24 hours later, the cells were labelled for 1 h at 37 °C and 5% CO_2_ atmosphere with 1 µM (SNAP-ligand) self-labelled BG-JF646, derived from JF646-NS (Tocris, 6148) and BG-NH_2_ (New England Biolabs Inc., S9148S) (36) in Fluorobrite DMEM medium with 10% (v/v) FBS. After this they were washed extensive and incubated for another 30 min in ligand-free Fluorobrite DMEM medium with 10% (v/v) FBS.

150.000 qKO cells (stably expressing FLAG-Cldns) were seeded on a 12 mm, polycarbonate, 0.4 µm Millicell cell culture insert (Merck, PIHP01250) for TJ barrier measurements. Medium was refreshed every 2-3 days, until cells were confluent and measured 7 days later. On the same day, the qKO cells were seeded in a µ-Slide 8-well glass bottom dish (Ibidi, 80827) for immunofluorescence assays, and in a 6-well plate (Corning, 3516) for immunoblotting. Medium was refreshed every 2-3 days. Fixation and lysis were performed 3-5 days after, when cells had reached confluency.

### Creation of stable cell lines

For the production of retrovirus, HEK293T cells were seeded in a 10 cm dish (Sarstedt, 833902), so they would be at roughly 50% confluency 24 hours later. The cells were then transfected with the calcium phosphate method. 15 μg of pLIB-CMV-FLAG-Cldn-Puro plasmid was combined with 10.5 μg pCIG3.NB packaging plasmid, 4.5 μg pMD2.G retroviral envelope plasmid, 60 μL 2M CaCl_2_ solution and 440 μL TE buffer (10 mM Tris, 2 mM EDTA, pH 8.0), and incubated for 5 min at RT. This solution was dropwise added into a 500 μL 2x HBS solution (50 mM HEPES, 280 mM NaCl, 1.5 mM Na_2_HPO_4_, pH 7.05) under slight agitation. After another 20 min incubation at RT, the resulting solution was used for transfection. After another 24 hours, the HEK293T medium was refreshed. Finally, virus-containing supernatant was collected by spinning down the medium for 5 min at 720 g at 3-4 and 5-6 days after transfection.

To generate stable cell lines, qKO cells were seeded so they would be at 50-60% confluency 48 hours later. The cells were then incubated with 4 mL virus-containing supernatant, filled up to 10 mL with normal medium. After 2-3 days of incubation, successfully transduced cells were selected through the addition of 2 μg/mL puromycin to the medium. Selection medium was refreshed every 2-3 days and cells were split when they were 80-90% confluent. After selection, cells were cultured to at least 50% confluency before using them for experiments.

### Fixation and immunocytochemistry

COS-7 cells for fixed STED imaging were fixed for 20 minutes at 37 °C with pre-warmed 4% (w/v) PFA and 4% (w/v) sucrose in PBS. qKO cells were fixed with pre-cooled 100% ethanol for 15 minutes at -20 °C and blocked for 30 minutes at RT with blocking buffer (6% (v/v) normal goat serum, 1% (w/v) BSA, 0.05% (v/v) Tween-20 in PBS). Primary antibody labeling was done for 2 hours at RT in blocking buffer. Rabbit anti-ZO1 antibody (1:100, Thermo Fisher, 617300) and mouse anti-FLAG antibody (1:200, Sigma-Aldrich, F3165) were used. After thorough washing with PBS, secondary antibody labeling in blocking buffer was done for 30 minutes at RT. For this, goat anti-rabbit Alexa 647 (1:200, Invitrogen, A21244) and goat anti-mouse Alexa 488 (1:200, Invitrogen, A11029) antibodies were used. Finally, cell nuclei were labelled with DAPI (1:5000, Thermo Fisher, 62248) and cells were once more washed thoroughly with PBS.

### Stimulated Emission Depletion (STED) microscopy

STED imaging was done with a STEDYCON system (Abberior Instruments GmbH, Göttingen), mounted on a Nikon Eclipse TI research microscope, equipped with a Plan APO Lambda 100x/1.45 NA lambda oil objective (Nikon) and controlled by NIS Elements (Nikon). 24 h prior to live-cell imaging, an incubation chamber surrounding the microscope set-up was set to 37 °C to provide stable focus during imaging. Imaging was performed in PBS (fixed cells) or in FluoroBrite DMEM medium with 10% (v/v) FBS (living cells) at 37 °C. Excitation was done with a 640 nm diode laser, STED depletion with a 775 nm laser. Emission was detected with a single counting avalanche photodiode (650-700 nm). A pixel size of 20 nm by 20 nm and a line average of 2 for living and 25 for fixed cells were used. For corresponding confocal images, the pixel size was 100×100 nm and no line averaging as performed. For time-lapse imaging, the acquisition time per frame was set to 1.96 seconds/frame.

### Confocal microscopy

Confocal images of qKO cells were acquired with an LSM780 equipped with spectral detector and PMTs from Carl Zeiss Microscopy. The LSM780 was controlled by the Zeiss Zen Black software (Carl Zeiss Microscopy). For determination of protein localization, multi-color confocal imaging was performed in sequential mode with the following fluorophore-specific excitation (Ex.) and emission filter (EmF.) settings: DAPI (Ex.: 405 nm; EmF.: 415–480 nm), Alexa 488 (Ex.: 488 nm; EmF.: 491–561 nm), Alexa 647 (Ex.: 633 nm; EmF.: 638–752 nm). A PL APO DIC M27 40 x/1.3 NA oil objective, and a zoom factor of 1 were used. Stacks of eight 512×512 pixel images at a distance of 1 µm were acquired, using a line average of 2. For the analysis of TJ ruffles, Alexa 488 (Ex.: 488 nm; EmF.: 491-578nm) and Alexa 647 (Ex.:633 nm; EmF.: 638-735nm) were sequentially imaged. Images were acquired with a PL APO DIC M27 63×/1.40 NA oil objective (Carl Zeiss Microscopy) at a zoom factor of 3 and a line average of 2. Stacks of 3-6 images (1024×1024 pixels) at 1 µm were obtained to cover the TJ in all cells in view.

### Image analysis

Image preparation and analysis was done in Fiji Is Just ImageJ (63).

### Meshwork classification

For 1 hour per condition, each COS-7 cell-cell overlap was systematically imaged. Subsequent classification into membrane staining, isolated strands, thin meshworks and dense meshworks was done independently by 4 individuals. Per cell-cell overlap on classifier was assigned, based on the majority vote during individual classification.

### Automated zigzag index quantification

To quantify the zigzag indexes in the ZO1 channel, the ‘Triangle Dark’ auto threshold was applied to them and they were converted to a mask. The mask was then dilated, smoothened with a median filter (radius: 12 pixels) and finally skeletonized. The resulting skeleton was analyzed with the Analyze Skeleton (2D/3D) function in Fiji (64). The resulting data were further analyzed in RStudio (version 4.0.3). First, any identified branches under 2 µm were eliminated, to remove anything outside of the cell-cell contacts. For all of the remaining branches, now considered cell-cell contacts, the zigzag index was quantified by division of the branch length (α) by the Euclidean distance (β) (Fig. 4*C*). Finally, the mean zigzag index of all cell-cell contacts in the image (ranging from 24 to 71) was calculated.

### Cell lysis

Cells were washed once with PBS and lysed with 100 µL ice-cold lysis buffer (1% Triton X-100, 20 mM HEPES, pH 7.4, 130 mM NaCl, 10 mM NaF, 0.03% PIC) on ice. After incubation on ice under constant agitation for 30 minutes, the lysates were centrifuged at 17,000 g for 20 min at 4 °C. The supernatant was collected and 1 µl it was combined with 499 µl H_2_O and 500 µl 2x Bradford reagent. After a 5-minute incubation the protein concentration was determined through measurement of the OD595 with a photometer (BioPhotometer plus, Eppendorf). Finally, 6x SDS sample buffer (0.375 M Tris-HCl (pH 6.8), 10% (w/v) SDS, 60% (v/v) glycerol, 0.6 M DTT, 0.06% (w/v) bromophenol blue) was added and protein lysates were denatured for 5 minutes at 95 °C.

### Immunoblotting

Protein size separation of 25 µg of protein was done on a NuPAGE 4-12% Bis-Tris gel (Invitrogen, NP0336BOX) in NuPAGE MES SDS-buffer (Invitrogen, NP0002) at 100 V for 120 minutes. The proteins were transferred to a nitrocellulose membrane (Cytiva, 1060004) in transfer buffer (10% (v/v) methanol, 25 mM Tris-HCl (pH 7.6), 192 mM glycin) for 90 minutes at 110 V on ice. The membrane was blocked with 5% (w/v) milk in TBS-T (0.1% (v/v) Tween 20, 0.1 M Tris-Base, 0.7 M NaCl, pH 7.6) for 1 hour at RT and subsequently incubated with primary antibody in 3% (w/v) BSA in TBS-T overnight at 4 °C. Primary antibodies used were mouse anti-FLAG antibody (1:1000, Sigma-Aldrich, F3165), mouse anti-β-actin antibody (1:10000, Sigma-Aldrich, A5441), rabbit anti-Cldn3 antibody (1:500, Invitrogen, 341700) and rabbit anti-Cldn4 antibody (1:500, Invitrogen, 364800). After extensive washing with TBS-T, the membrane was incubated for 1 hour at RT with secondary antibody in 5% (w/v) milk in TBS-T. Secondary antibodies used were goat anti-mouse HRP (1:5000, Jackson, 115035003) and goat anti-rabbit HRP (1:5000, Jackson, 111035003). After another round of extensive washing with TBS-T and TBS (0.1 M Tris Base, 0.7 M NaCl, pH 7.6), proteins were visualized by the addition of HRP substrate (SuperSignal™ West Pico PLUS Chemiluminescent Substrate, Thermo Fisher, 34580) and imaging with a ChemiDoc XRS+ imaging system (BioRad) controlled by the Image Lab software (version 6.0.1).

### Transepithelial electrical resistance (TER) and fluorescein permeability measurements

TER and fluorescein flux measurements were done in an Ussing-Chamber that has been designed to fit the Millicel filters (65). TER was measured with 5 mL of Ringer’s solution (1.2 mM CaCl_2_, 10 mM glucose, 3 mM HEPES, 5.4 mM KCl, 1 mM MgSO_4_, 119 mM NaCl, 21 mM NaHCO_3_; pH 7.4) on each side. The values were corrected for the resistance of the filter holders and solution on their own. Through constant bubbling (with 95% O_2_ and 5% CO_2_) and heating (to 37 °C), the Ringer’s solution was kept in equilibrium. Before measuring the fluorescein flux, the Ringer’s solution on both sides was increased to 10 mL. After application of a voltage clamp, 10 µl of 100 mM fluorescein was added on the apical side. For 15 minutes a small sample was collected in a 96 well plate (Corning, 3365) from the basolateral side every 5 minutes, and replenished with Ringer’s solution. Fluorescein concentrations were determined with a Tecan Inifite 200 plate reader (Tecan Trading AG, Männedorf, Switzerland) through excitation at 490 nm and detection of emission at 525 nm.

### Sequence alignments

In this paper we mostly adhere to the Cldn nomenclature as suggested by Mineta et al. (7) (Table 1). Cldn27 as suggested in this paper was excluded, as this protein has been shown to be distinct from the Cldn family (8). All Cldn sequences were aligned in Clustal Omega (EBML-EBI) (66) and the outcome was presented as a phylogenetic tree with Drawtree (3.67, phylogeny.fr) (67). For the specific alignment of Cldn3, Cldn4, Cldn6, Cldn8, Cldn9 and Cldn17, PSI/TM-Coffee (version 11.00) was used, with the ‘transmembrane’ and ‘UniRef100’ homology search options (68).

**Table 1.**
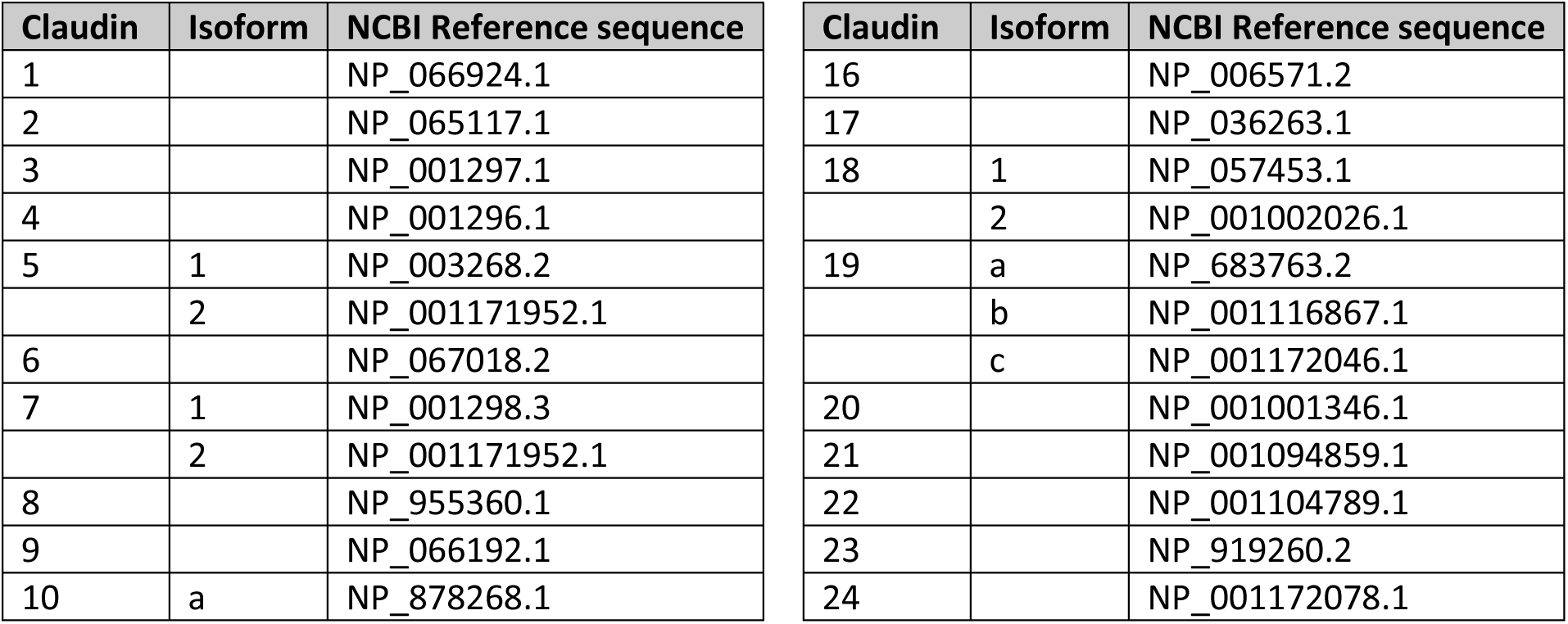
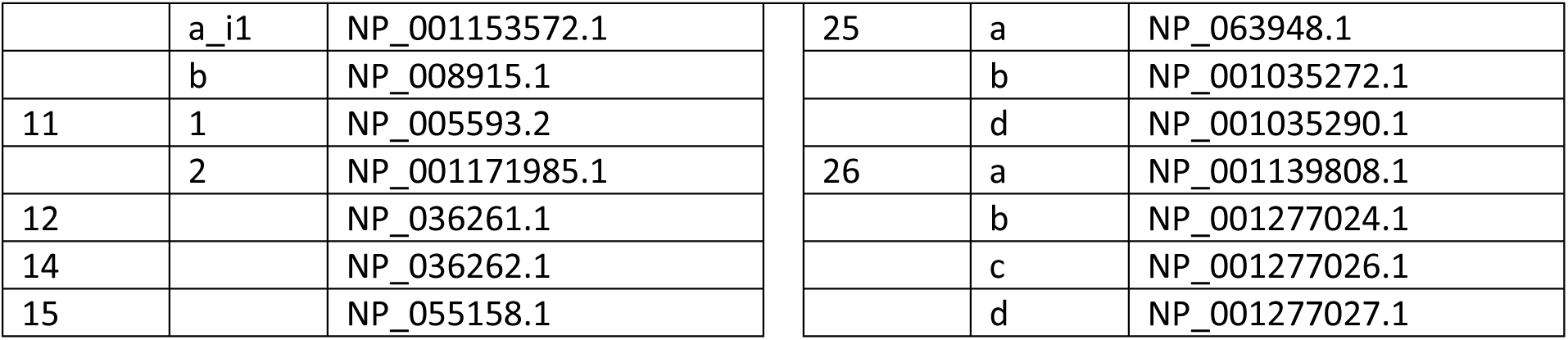
NCBI Reference sequences used in this study.

| Claudin | Isoform | NCBI Reference sequence | Claudin | Isoform | NCBI Reference sequence |
| --- | --- | --- | --- | --- | --- |
| 1 |  | NP_066924.1 | 16 |  | NP_006571.2 |
| 2 |  | NP_065117.1 | 17 |  | NP_036263.1 |
| 3 |  | NP_001297.1 | 18 | 1 | NP_057453.1 |
| 4 |  | NP_001296.1 |  | 2 | NP_001002026.1 |
| 5 | 1 | NP_003268.2 | 19 | a | NP_683763.2 |
|  | 2 | NP_001171952.1 |  | b | NP_001116867.1 |
| 6 |  | NP_067018.2 |  | c | NP_001172046.1 |
| 7 | 1 | NP_001298.3 | 20 |  | NP_001001346.1 |
|  | 2 | NP_001171952.1 | 21 |  | NP_001094859.1 |
| 8 |  | NP_955360.1 | 22 |  | NP_001104789.1 |
| 9 |  | NP_066192.1 | 23 |  | NP_919260.2 |
| 10 | a | NP_878268.1 | 24 |  | NP_001172078.1 |

|  |  |  |  |  |  |  |
| --- | --- | --- | --- | --- | --- | --- |
|  | a_i1 | NP_001153572.1 |  | 25 | a | NP_063948.1 |
|  | b | NP_008915.1 |  |  | b | NP_001035272.1 |
| 11 | 1 | NP_005593.2 |  |  | d | NP_001035290.1 |
|  | 2 | NP_001171985.1 |  | 26 | a | NP_001139808.1 |
| 12 |  | NP_036261.1 |  |  | b | NP_001277024.1 |
| 14 |  | NP_036262.1 |  |  | c | NP_001277026.1 |
| 15 |  | NP_055158.1 |  |  | d | NP_001277027.1 |

### Structural modeling and molecular dynamics simulation

Cldn monomers and oligomers were handled, modeled, simulated und visualized using the Schrödinger Maestro BioLuminate software platform (BioLuminate, version 4.9.134, Release 2022-4, Schrödinger, LLC, Germany, 2022), Schrödinger PyMOL 2.5.2 (http://www.pymol.org/pymol) and Linux-x86_64 GPU computing workstations. Cldn3 homology oligomer models were generated using the murine sequence 1-187 (Uniprot ID Q9Z0G9) and the membrane-equilibrated dodecamer model of human Cldn10b (8IBli; (29)) as template. Using the PyMod 3.0.2. plugin for PyMol (http://schubert.bio.uniroma1.it/pymod; (69)), each of the 12 Cldn chains/subunits of the Cldn10b dodecamer was replaced by the corresponding Cldn3 homology model. The dodecamer was minimized using the ‘MacroModel Minimization’ module of BioLuminate and a water-solvated environment with a gradient convergence threshold of 0.05 kJ mol−1 Å−1. This dodecamer model consisted of two trans-interacting cis-hexamers resulting in three interlocked neighboring cis-/trans-tetrameric building blocks. Dodecamers were modeled since the middle building blocks are fully connected to the adjacent ones, similar as proposed for polymeric Cldn strands (29). The MD simulations were carried out using the ‘Desmond Molecular Dynamics’ module of BioLuminate (D. E. Shaw Research, New York, NY, 2021. Maestro-Desmond Interoperability Tools, Schrödinger, New York, NY, 2021; (29)). Systems were generated using TIP3P waters, charge-neutralizing ions and 0.15 M NaCl, and the simulations were performed with an OPLS3e force field in NPT ensemble, a temperature of 310 K and a pressure of 1.01325 bar (29). ‘Desmond Minimization’ was performed for 100 ps and the systems were relaxed using BioLuminate default protocol. Afterwards, the Cldn3 system was equilibrated stepwise by lowering the constraints on the protein ECS backbone and sidechains from 5 to 0 kcal mol-1 Å-² (force constant) over 80 ns while keeping the constraints for the backbone of transmembrane helices on 5 kcal mol-1 Å-². After equilibration, the Cldn3 model was simulated with only the transmembrane helices constrained (3 kcal mol-1 Å-²) for 50 ns (production run MD1). In addition, a stepwise equilibration was performed for 20 ns and followed by simulation with 5 kcal mol-1 Å-² only on the backbone of transmembrane helices for 50 ns (production run MD2). Salt interactions between E153 and R157 were analyzed using the MD trajectories and analysis tools within Bioluminate.

### Statistics

Data visualization and statistical analyses were performed in GraphPad Prism (version 9.5.1). Gaussian distribution of the data was verified with multiple normality tests (D’Agostino & Person test, Anderson-Darling test and Shapiro-Wilk test).

## Supporting information

Video 1. Live-STED imaging of a SNAP-Cldn4 meshwork between two COS-7 cells. Scale bar: 1 micrometer

## Supporting information

**Supplementary figure 1.**
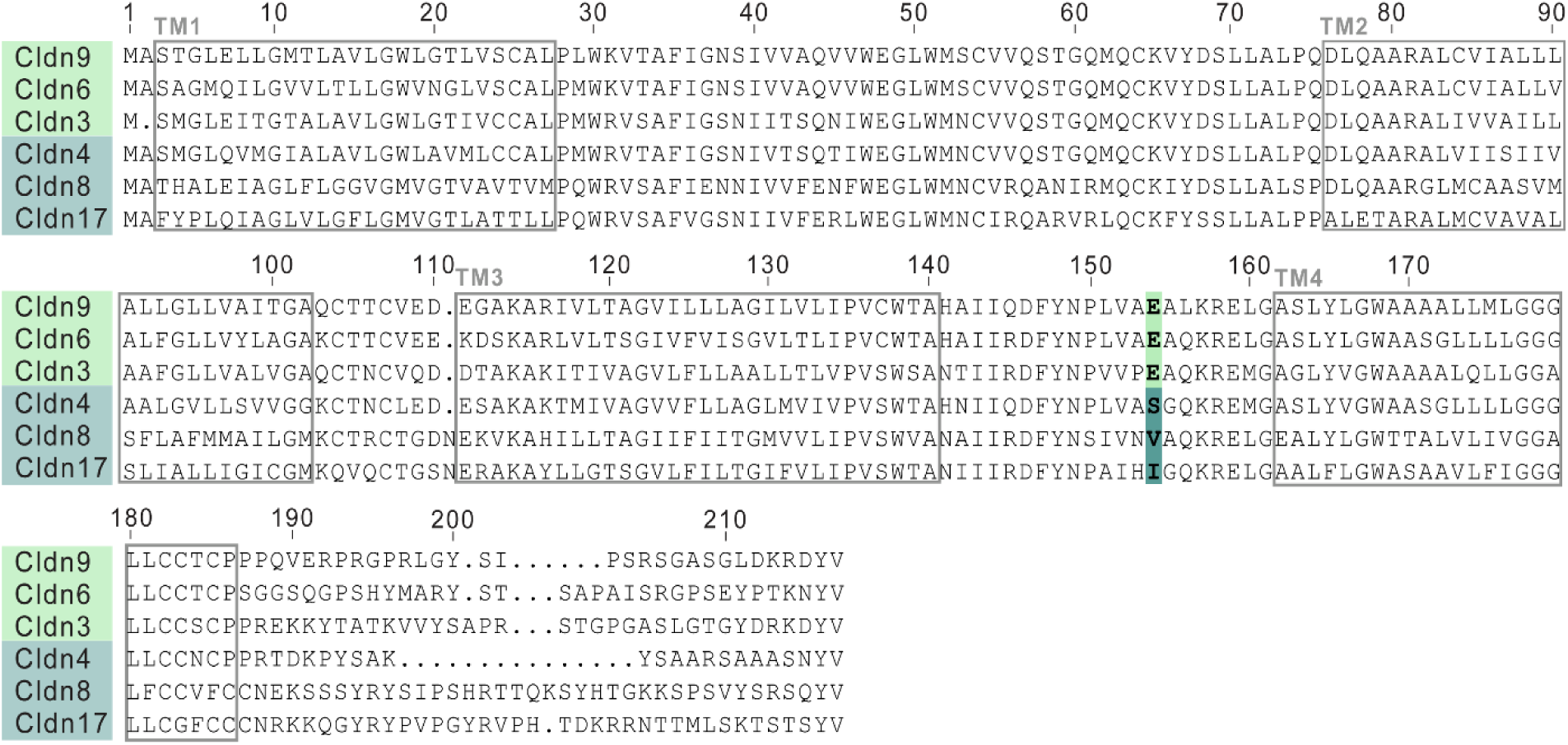
Sequence alignment of Cldn3, Cldn4, Cldn6, Cldn8, Cldn9 and Cldn17 demonstrates a key difference between meshwork versus non-meshwork formers. Meshwork formers are indicated in green, non-meshwork formers in blue. Human sequences were used and the numbering is according to Cldn9. The transmembrane helices are indicated in grey and the identified difference (position 153 in ECS2) is highlighted.

